# CAGEE: computational analysis of gene expression evolution

**DOI:** 10.1101/2022.11.18.517074

**Authors:** Jason Bertram, Ben Fulton, Jason P. Tourigny, Yadira Peña-Garcia, Leonie C. Moyle, Matthew W. Hahn

## Abstract

Despite the increasing abundance of whole transcriptome data, few methods are available to analyze global gene expression across phylogenies. Here, we present a new software package (CAGEE) for inferring patterns of increases and decreases in gene expression across a phylogenetic tree, as well as the rate at which these changes occur. In contrast to previous methods that treat each gene independently, CAGEE can calculate genome-wide rates of gene expression, along with ancestral states for each gene. The statistical approach developed here makes it possible to infer lineage-specific shifts in rates of evolution across the genome, in addition to possible differences in rates among multiple tissues sampled from the same species. We demonstrate the accuracy and robustness of our method on simulated data, and apply it to a dataset of ovule gene expression collected from multiple self-compatible and self-incompatible species in the genus *Solanum* to test hypotheses about the evolutionary forces acting during mating system shifts. These comparisons allow us to highlight the power of CAGEE, demonstrating its utility for use in any empirical system and for the analysis of most morphological traits. Our software is available at https://github.com/hahnlab/CAGEE/.

## Introduction

Early studies of gene expression in single genes revealed widespread and frequent changes in the levels, timing, and breadth of expression across species (reviewed in Wray et al. 2003; Fay and Wittkopp 2008; Hill et al. 2021). Such changes in gene expression have been shown to be responsible for many differences between species, and may be a major driver of evolution (King and Wilson 1975). Advances in sequencing technologies (i.e. RNA-seq) have transformed research into gene expression, allowing researchers to cheaply and accurately measure transcript levels for every gene in a genome, in multiple tissues, and across several timepoints or conditions (Wang et al. 2009). There is now a flood of interest in applying RNA-seq to whole clades of organisms in order to identify the genetic changes and evolutionary forces driving species differences (e.g. Brawand et al. 2011; Meisel et al. 2012; Coolon et al. 2014; Harrison et al. 2015; Berthelot et al. 2018; Catalan et al. 2019; Blake et al. 2020; El Taher et al. 2021).

To better understand the importance of changes in gene expression, researchers must be able to characterize the mechanisms and modes by which gene expression evolves. Such work entails understanding the role of natural selection in driving species differences, the stages of development or the tissues that evolve most rapidly, as well as the environments most likely to generate changes in gene expression (Dunn et al. 2013; Hill et al. 2021; Price et al. 2022). Phylogenetic comparative methods enable the rigorous study of traits like gene expression across a species tree (Revell and Harmon 2022). These methods can be used for testing hypotheses about natural selection, the inference of ancestral states (allowing us to polarize the direction of changes), and the estimation of evolutionary rates. Multiple software packages are available that implement a wide variety of comparative methods (e.g. Pennell et al. 2014), including models specifically intended for studying gene expression across a tree (Bedford and Hartl 2009; Rohlfs et al. 2014; Rohlfs and Nielsen 2015; Catalán et al. 2019; Chen et al. 2019; Yang et al. 2019).

However, as far as we are aware, all existing comparative methods for analyzing gene expression implement fundamentally single-gene analyses. Each gene is considered a separate trait, such the evolutionary parameters for each gene are estimated separately. Single-gene analyses can be used to identify tissue-specific or lineage-specific shifts in evolutionary rates, but their power is quite low (Beaulieu et al. 2012). As a result, identifying trends in evolution must be carried out *post hoc* by summing the number of genes found to be individually significant (e.g. Harrison et al. 2015; El Taher et al. 2021). This approach is less than ideal, especially when carrying out comparisons between branches of different lengths or between tissues with different average expression levels (both of which can result in differential statistical power).

Therefore, to better characterize the forces affecting gene expression evolution, we must be able to model effects shared along a lineage, experienced by many genes in the same tissue, or experienced by all genes found in the same environment. In this article, we present a genome-scale platform for the analysis of gene expression data that allows for such shared factors. Our software, CAGEE (Computational Analysis of Gene Expression Evolution), provides a robust set of methods for analyzing expression data across a species tree. CAGEE estimates ancestral states and rates, with rates shared by all or subsets of genes (single-gene analyses can also be carried out). We show that lineage-specific and tissue-specific (or condition-specific) rates can be accurately inferred, and provide principled statistical approaches for model selection. Our current implementation uses a bounded Brownian motion model and assumes expression data are accurate, but the architecture and codebase will easily allow for future extensions that relax these and other assumptions.

### New Approaches

We model gene expression evolution as a bounded Brownian motion (BBM) process on a known species tree (cf. Boucher and Démery 2016). Our model has a single bound: trait values must be greater than or equal to zero; there is no upper bound (Figure 1). Previous researchers have often modeled gene expression using an Ornstein-Uhlenbeck (OU) process (e.g. Bedford and Hartl 2009; Rohlfs et al. 2014; Rohlfs and Nielsen 2015; Chen et al. 2019), a model that includes a force constraining traits about the mean. However, to our knowledge, the OU model has only been compared against an unbounded Brownian motion model (i.e. one that allows negative expression values), making fair comparisons difficult. In addition, OU models may be frequently and incorrectly favored over simpler models due to several biases (e.g. measurement error), especially when the number of tips in a tree is small (Pennell et al. 2015; Silvestro et al. 2015; Boucher and Démery 2016; Cooper et al. 2016; Catalán et al. 2019). Therefore, the initial version of our software models gene expression with the BBM process, which naturally bounds possible values without invoking an additional constraining force.

**Figure 1.**
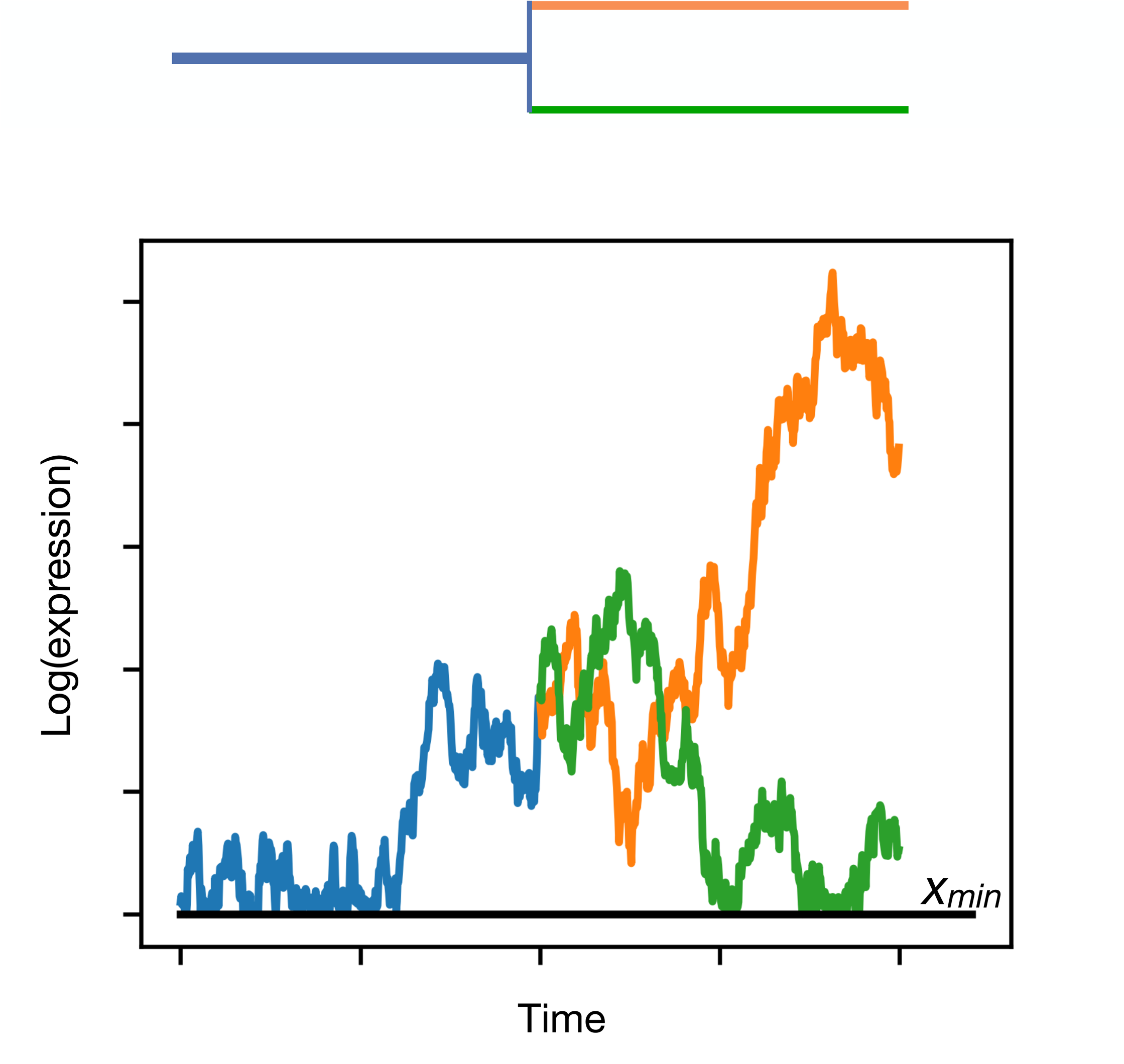
Bounded Brownian motion model. An example trait is shown in the bottom graph, evolving along the tree shown above. Although the data input to CAGEE are linear expression levels, internally it logs expression to ensure higher variance among more highly expressed genes. There is also a minimum value, *xmin*, added to all tips.

Let 𝐸_ij_ ≥ 0 be the expression level of gene 𝑖 in species 𝑗. We assume that log-transformed expression 𝑋_ij_ = ln(𝐸_ij_ + 𝑒_m$n_) evolves as a Brownian motion process with variance 𝜎^2^ per unit time, where 𝑒_m$n_ is a small offset (constant across genes and species) that prevents 𝑋_ij_ from taking infinite values if measured values of 𝐸_ij_ are zero. We log-transform before assuming Brownian motion because we expect the variance in the evolutionary process to scale with expression level. Assuming that 𝐸_ij_ is itself Brownian would unrealistically assume that the rate of evolution is constant across expression levels, even though expression levels vary by many orders of magnitude. We impose a reflecting lower boundary at 𝑥_m$n_ = ln(𝑒_m$n_), meaning that the Brownian walk immediately bounces back if it reaches 𝑥_m$n_. Expression can therefore effectively never reach zero, our theoretical lower bound (Figure 1).

The second major feature of our model (as implemented in CAGEE) is that many genes can share the evolutionary rate parameter, 𝜎^2^. This rate may be shared among genes expressed in the same tissue or sample, among genes located on the same chromosome, or among genes evolving along the same lineage of the phylogenetic tree. The simplest model allows 𝜎^2^ to be shared among all genes, providing an average rate of evolution across the genome and over time; this average may include genes that vary in their individual rates of evolution. We explain this model briefly here, with more detail provided in the Materials and Methods.

CAGEE infers the most likely value(s) of 𝜎^2^ consistent with an ultrametric tree, 𝑇, and a set 𝐸_{ij}_ of measured expression values at the tips of the tree; i.e. it maximizes the likelihood 𝐿(𝜎^2^|𝐸_{ij}_, 𝑇). Each gene is assumed to evolve independently, and so the likelihood for each gene 𝐿_i_(𝜎^2^|𝐸_i{j}_, 𝑇) is computed independently. The overall likelihood is obtained as the product 𝐿5𝜎^2^6𝐸_{ij}_, 𝑇7 = Π_i_𝐿_i_(𝜎^2^|𝐸_i{j}_, 𝑇) across genes. The likelihood for each gene 𝐿_i_(𝜎^2^|𝐸_i{j}_, 𝑇) is computed using the pruning algorithm (Felsenstein 1973). The key ingredient needed to apply the pruning algorithm is the transition probability density 𝑝5𝑥_𝑡_6𝑥_𝑡0_7 = Pr [𝑋(𝑡) = 𝑥_𝑡_|𝑋(𝑡_*_) = 𝑥_𝑡0_] for log-expression at time 𝑡 conditional on having log-expression 𝑥_𝑡0_ at time 𝑡_*_ along a lineage. CAGEE computes the transition density by solving the standard Brownian diffusion equation with reflecting boundary conditions (Materials and Methods). The transition density is used to propagate expression probabilities along the tree: if the probability density of log-expression at time 𝑡_*_ is 𝑓(𝑥_𝑡0_), then the probability density at time 𝑡 on the same lineage is 𝑓(𝑥_𝑡_) = ∫ 𝑝5𝑥_𝑡_6𝑥_𝑡0_7𝑓5𝑥_𝑡0_7𝑑𝑥_𝑡0_. At each tip the probability density 𝑓(𝑥_𝑡0_) is a delta function centered at the corresponding measured value of 𝑋_ij_.

Starting with the known tip distributions, the pruning algorithm propagates back to the tips’ parent nodes. The distribution at the parent node is then the product of the two backward-propagated child node distributions. Proceeding iteratively across the tree, we ultimately obtain the gene-specific probability density for expression value at the root 𝑓_i_(𝑥_𝑅_). Viewed as a likelihood for 𝜎^2^, 𝑓_i_(𝑥_𝑅_) is the gene-specific likelihood conditional on the unknown ancestral root value; i.e. 𝑓_i_(𝑥_𝑅_) = 𝐿_i_(𝜎^2^|𝐸_i{j}_, 𝑇, 𝑥_𝑅_).

Therefore, we integrate over all possible 𝑥_𝑅_ to obtain,

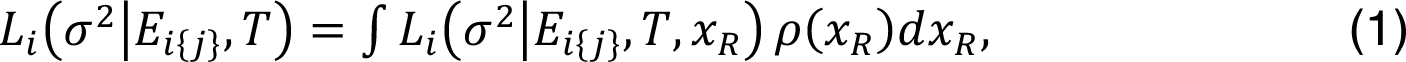

where 𝜌(𝑥_𝑅_) is the prior distribution for the root value of a randomly selected gene.

The default prior 𝜌(𝑥_𝑅_) is assumed to be a gamma distribution with 𝑘 = 0.375 and 𝜃 = 1600, though this distribution can also be set by the user in CAGEE. This choice is based on estimated expression distributions across genes in individual species, which we take as our baseline for the ancestral distribution. CAGEE uses the Nelder-Mead simplex method to find the optimal value(s) of 𝜎^2^.

## Results

### Using CAGEE

The required inputs for CAGEE are a Newick-formatted, rooted, ultrametric tree (with branch lengths) and a tab-delimited data file containing the expression levels of all species or taxa being studied. The data file can consist of data on one gene/transcript or thousands of different genes. The first line of the data file should contain the species’ names (matching those used in the Newick tree). In addition, headers for gene names, gene descriptions, and sample IDs (see next section for an explanation of “samples” in CAGEE) can be used. Subsequent lines each correspond to a single gene and contain expression levels for each species. Missing data can be denoted using multiple characters (-/?/N). Examples of Newick trees and corresponding data files can be found in the online user manual (https://github.com/hahnlab/CAGEE/docs/manual/cagee_manual.md).

We expect that CAGEE will most often be used to calculate the following outputs: one or more 𝜎^2^ values, ancestral states at each internal node (including 95% credible intervals around these states), and the final likelihood associated with a model. However, users do not have to search for 𝜎^2^: if a value for this parameter is specified, then the output of CAGEE will just be the ancestral states and a likelihood. In addition to the raw outputs provided in multiple formats (both tab-delimited files and NEXUS-formatted files), CAGEE computes basic statistics about changes in expression levels by comparing values at parent and child nodes. Summaries of these inferred changes for every gene and for every branch of the tree are output, so that the evolutionary history of gene expression changes in every gene are accessible to users. To avoid over-interpretation of small changes in inferred expression levels—especially when there is uncertainty in ancestral states—CAGEE will also compare the credible intervals at parent and child nodes to note if a change is “credible” (i.e. the intervals do not overlap). Credible intervals are calculated by summing the probabilities across possible ancestral states at each node, so that 95% of the probability density is included. Credible changes on each branch are annotated as such in the output.

We most often expect that an ultrametric species tree will be used as the input topology, but this is not required by CAGEE. If users wish to specify a gene tree, or some other bifurcating tree, as input, those can be used in CAGEE as well. However, the major advantage of CAGEE—incorporating information from multiple genes to accurately estimate genome-wide rates—will rapidly diminish for trees that represent the history of only a minority of the genome. Trees that include duplication events should provide suitable estimates for any genes that follow this topology, but CAGEE does not have a way to further combine disparate gene trees.

There are multiple options available for running CAGEE. Users who can take advantage of multiple threads can specify the number to use on the command line. Complex models can also take a long time to converge; by default, CAGEE runs a maximum of 300 iterations of the Nelder-Mead search, but users can increase this number in subsequent runs if the likelihood is still improving when the limit is hit. As mentioned above, the default prior distribution for the root state is a gamma distribution with 𝑘 = 0.375 and 𝜃 = 1600. This distribution can also be specified by the user if desired. Information on how to run more complex evolutionary models, beyond a single 𝜎^2^, is given in the next section.

### Estimating evolutionary rates in CAGEE

We tested CAGEE’s ability to accurately estimate 𝜎^2^ by varying this rate parameter and the number of genes used for inference, as well as the amount of missing data in each dataset. We simulated different single values of 𝜎^2^ across a tree with constant branch lengths (Supplementary Figure 1) using the simulation tool available within CAGEE. (Note that the total amount of evolution in a tree is determined by the product 𝜎^2^ ⋅ 𝑡, such that changes in branch lengths will have an effect commensurate with changes in 𝜎^2^.) Figure 2 shows the average error associated with estimates of different 𝜎^2^ values and using different numbers of genes within each dataset. As can be seen, the error across all parameter values and dataset sizes is quite small (generally less than 2.5%), and is less variable for larger dataset sizes. Fortunately, we expect that most empirical datasets will contain closer to 10,000 genes than 1,000 genes. The results in Figure 2 are for an ancestral state vector of length *N*=200 (the default setting in CAGEE; Materials and Methods); we also estimated 𝜎^2^ when allowing the ancestral state vector to have length *N*=500 (Supplementary Figure 2A). There appears to be minimal gain from increasing the resolution in this vector, though the computational time is greatly increased (similar to results in Boucher and Démery 2016). We evaluated the accuracy of CAGEE when different amounts of data were randomly missing: from 0% to 75% for a dataset of 1,000 genes. As shown in Supplementary Figure 2B, CAGEE has high accuracy even when large amounts of data are missing (at random) from a dataset.

**Figure 2.**
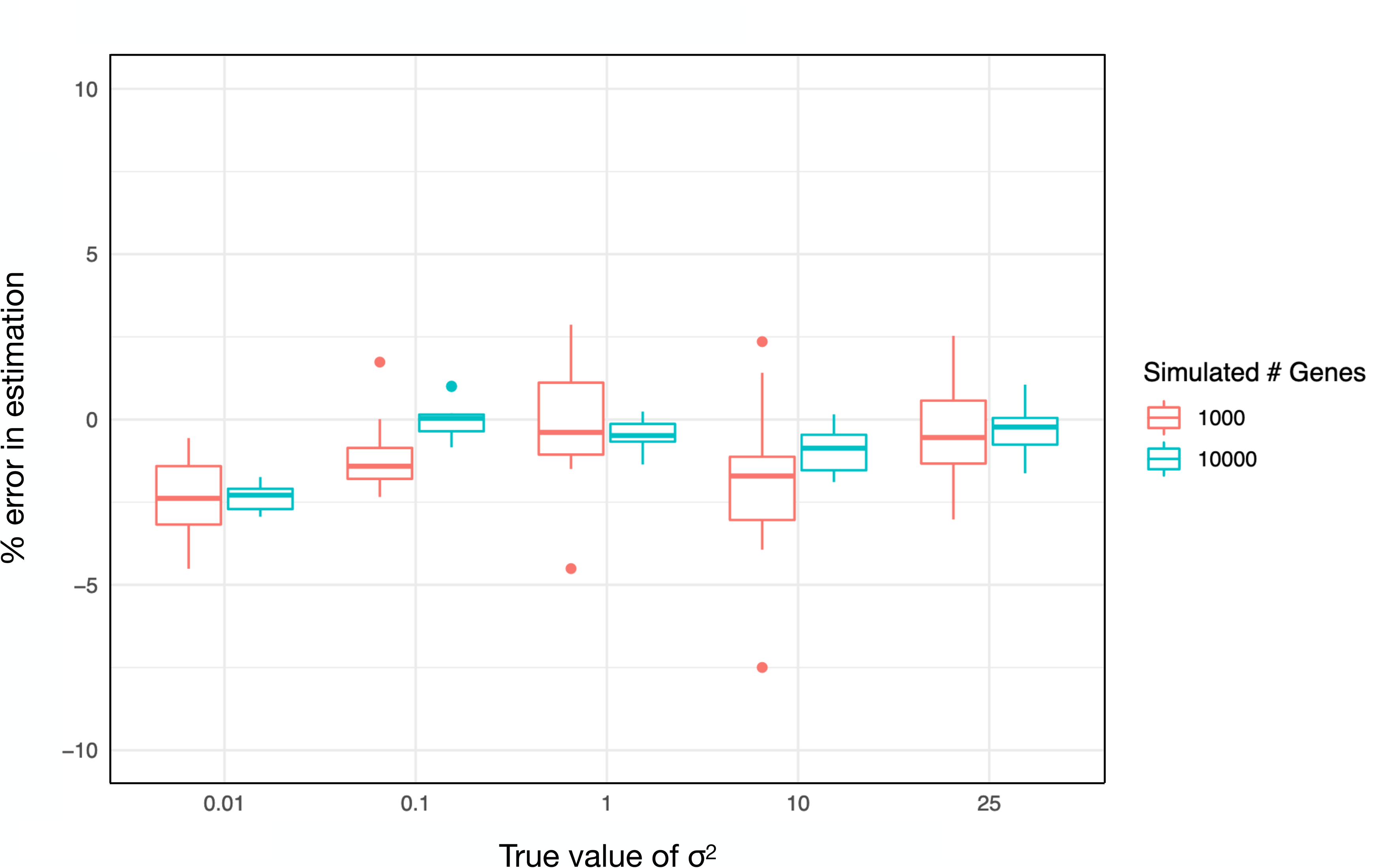
Accuracy of CAGEE. For five different values of 𝜎^2^ we simulated 1000 datasets, with each dataset comprised of either 1000 genes or 10000 genes. All genes in a dataset shared the same 𝜎^2^, but their values at the root were drawn independently from the prior. We then provided each simulated dataset to CAGEE in order to infer 𝜎^2^. Each box-and-whisker plot shows the mean (horizontal line), 50% interquartile range (box), 1.5X the interquartile range (vertical lines), and outliers (dots).

One major advantage of using CAGEE is that it combines information from multiple genes to infer a rate of evolution: this is why it can return estimates with high accuracy even when a large fraction of the data are missing. To further demonstrate this advantage, we simulated evolution in 1,000 genes using the same parameter value (𝜎^2^=1) and then estimated 𝜎^2^ for each of the 1,000 genes individually. Supplementary Figure 2C shows that these individual estimates of 𝜎^2^ are quite error-prone: although the mean of all genes is close to the true value, individual estimates can be 7-8X higher or lower and there is a large amount of variance. Although we have not shown it here, we do expect that the accuracy of 𝜎^2^ will be greater for trees with larger numbers of tips, even for estimates derived from single genes (cf. O’Meara et al. 2006). On the other hand, CAGEE is combining information from multiple genes to infer an *average* rate of evolution, even when the underlying rate may be quite variable. To explore any effect of underlying rate variation, we carried out further simulations that combined three simulations of 1,000 genes each with 𝜎^2^ equal to 0.5, 3, and 9, respectively (we repeated these simulations 10 times). When analyzed as single datasets with 3,000 genes total, the average 𝜎^2^ inferred was 3.76, approximately 9% lower than the arithmetic mean rate (Supplementary Figure 2D). It is well-known that single-rate phylogenetic likelihood models tend to underestimate rates of evolution when there is underlying variation (Golding 1983; Gillespie 1986; Yang 1996; Mendes et al. 2020), and we see this effect here. Fortunately, the bias is small, and can be corrected in the future by including gamma-distributed rate variation into CAGEE. Overall, inferences of 𝜎^2^ should be quite accurate when a single rate parameter is shared across the tree and across all genes and lineages.

Variation in the rate of expression can currently be accommodated by CAGEE in a number of ways, using multi-rate 𝜎^2^ models. One type of model allows users to specify that their data come from different “samples”: these samples can represent tissues, conditions, timepoints, and even subsets of the genome (e.g. the X chromosome, or a specific functional class of genes). In the input data file, the “SAMPLETYPE” column is used to indicate which sample each gene is a member of; a separate 𝜎^2^ value will be calculated for each sample or set of samples (these values are assumed to be shared among all lineages in the tree). Specifying more than one sample means that an individual gene or transcript name can be used more than once (i.e. once for each sample), but there is no requirement that genes are measured in each sample. For instance, assigning all autosomal genes to sample 1 and all X-linked genes to sample 2 would not permit for any overlap in gene assignment, but is perfectly allowable in CAGEE.

Each additional sample requires another 𝜎^2^ parameter to be estimated, and often researchers would like to know if fitting this extra parameter is justified by the data. Under standard information-theoretic criteria (Burnham and Anderson 2002), twice the difference in log-likelihoods between nested models should be χ^2^-distributed with degrees of freedom equal to the difference in the number of parameters between models. To test this expectation, we simulated 1000 datasets with a single 𝜎^2^ value, but fit models with two 𝜎^2^ values (assigning 1000 genes to two equal-sized samples at random; the relative size of the samples should not affect the false positive rate). As anticipated, the results fit a χ^2^ distribution with one degree of freedom, with 4.4% of datasets having a difference in 2*log-likelihood greater than 3.84 (5% are expected by chance). This indicates that standard statistical procedures should adequately control the false positive rate when fitting multi-sample 𝜎^2^ models.

CAGEE also allows models in which 𝜎^2^ varies across branches of the species tree. It does so by fitting separate 𝜎^2^ parameters for different parts of the tree. On the command line, CAGEE enables users to specify how multiple 𝜎^2^ parameters should be assigned to branches. For *n* taxa, from 1 to 2*n*-2 parameters can be specified, and branches can be grouped together in any way. For instance, a two-parameter model can have all branches that share a rate adjacent to one another in the tree (Supplementary Figure 3A) or spread out across the tree (Supplementary Figure 3B). Similar to the analyses carried out above for the false positive rate associated with multiple samples, we simulated data with a single 𝜎^2^ value and then fit models with multiple 𝜎^2^ parameters. Regardless of how we distributed the two rate classes across the tree we observed good control of the false positive rate: 4.5% and 5.4% of 1000 simulated datasets were significant at the *P*=0.05 level (for the trees shown in Supplementary Figures 3A and 3B, respectively). More limited simulations also showed that we could accurately estimate multiple 𝜎^2^ parameters when the data were simulated with multiple rates (Supplementary Table 1). Together, our results suggest that we can estimate multiple types of multi-rate models, and can accurately control the false positive rate when doing so.

### Analysis of wild tomato transcriptome data

To demonstrate the utility of CAGEE in an empirical system, we analyzed data from a clade that includes domesticated tomato, *Solanum lycopersicum*. This dataset contains gene expression levels in unfertilized ovules from the flowers of six species, one of which (*S. pennellii*) has two different populations represented (Figure 3). There are 14,556 genes with expression levels measured in all seven accessions. RNA-seq data for five of the seven accessions have been published previously (Moyle et al. 2021; Hibbins and Hahn 2021), while two others are presented here for the first time (Materials and Methods). Note, however, that all data were collected from all samples at the same time (Materials and Methods).

**Figure 3.**
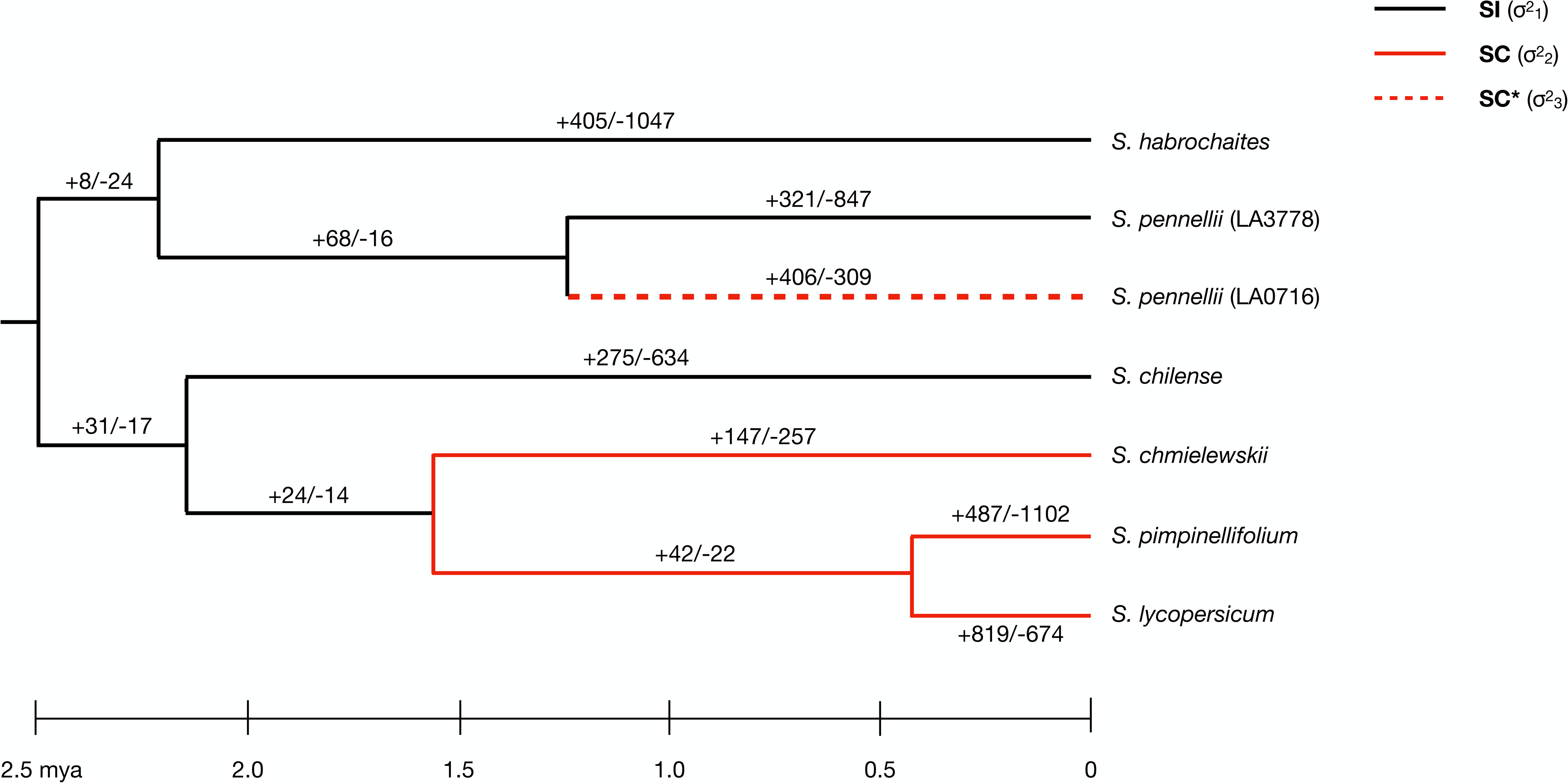
Changes in gene expression along the tomato phylogeny. Given the set of relationships among the seven *Solanum* accessions used here, we tested multiple models that had branches assigned as different 𝜎^2^ parameters (Table 1). In model A, all branches share 𝜎^2^_1_. In model B, all black branches share 𝜎^2^_1_, while all red branches share 𝜎^2^_2_. In model C, all black branches and the dashed red branch share 𝜎^2^𝜎^2^_1_, while all & 1 solid red branches share 𝜎^2^_2_. In model D, all black branches share 𝜎^2^, all solid red & 1 branches share 𝜎^2^, and the dashed red branch is assigned 𝜎^2^_3_. Using the results from & 3 model D, we inferred the number of genes that had credible increases or decreases in expression level along each branch (results for all changes are shown in Supplementary Figure 4). Numbers are reported as +increases/-decreases for each branch.

Most species within the tomato clade are self-incompatible (SI), the ancestral state in the family Solanaceae (Igić et al. 2006). Self-incompatibility means that plants must outcross in order to successfully fertilize ovules. However, self-compatibility (SC) has evolved multiple times both within the Solanaceae and within the genus *Solanum* (Goldberg et al. 2010; Bedinger et al. 2011). Self-compatible individuals are able to successfully fertilize ovules using their own pollen, though many also still outcross (Whitehead et al 2018; including in *Solanum*: Vosters et al. 2014 and references therein). Importantly, we have *a priori* expectations about the rate at which reproductive traits—including ovule gene expression—might evolve between groups with different mating systems. Due to conflict within and between the sexes, it is generally expected that reproductive traits in species that outcross more (i.e. SI taxa) should evolve more rapidly than in species that inbreed more (i.e. SC taxa; Clark et al. 2006). Such patterns are found in some analyses of the rate of protein evolution (e.g. Gossmann et al. 2016; Harrison et al. 2019), but are equivocal in other comparisons (e.g. Gossmann et al. 2014, Moyle et al. 2021). These complex patterns might reflect additional effects that also accompany mating system shifts; for instance, such shifts often lead to reductions in effective population size in more selfing lineages (Charlesworth and Wright 2001). Mating system shifts could also alter global patterns of molecular evolution (including gene expression) by changing the strength and pattern of purifying selection, as morphological changes often accompany mating system changes. The exact effect of shifts in mating system on molecular evolution remains an open question.

The *Solanum* species sampled here represent two independent transitions from SI to SC, with one of the transitions (in accession *S. pennellii* LA0716) occurring recently enough that different populations within this species have different incompatibility systems (Figure 3). We therefore fit a series of nested models within CAGEE to test two related hypotheses about ovule gene expression evolution. First, we would like to know whether the rate of evolution of ovule gene expression is different in SI species than in SC species. Second, given the recent transition to SC within accession *S. pennellii* LA0716, we wanted to know if it shows a pattern of evolution more similar to SI or to SC species. In total, we fit four separate evolutionary models (Table 1; Figure 3). Model A has a single rate parameter for the entire tree. Model B has two rate parameters, one for SI species and one for SC species. This model assigns the branch leading to *S. pennellii* LA0716 as SC. Model C also has two rate parameters, one for SI and one for SC, but assigns *S. pennellii* LA0716 as SI. Model D has three rate parameters: one for SI species, one for longer-term SC species, and one for *S. pennellii* LA0716.

**Table 1.** Model parameters estimated from the tomato data.

| Model | Number of rates | -lnL | $\sigma^2_1$ | $\sigma^2_2$ | $\sigma^2_3$ |
| --- | --- | --- | --- | --- | --- |
| A | 1 | 67252.4 | 0.102 |  |  |
| B | 2 | 65883.9 | 0.074 | 0.134 |  |
| C | 2 | 65124.5 | 0.075 | 0.152 |  |
| D | 3 | 65108.6 | 0.077 | 0.152 | 0.067 |

Estimated results from the different models are shown in Table 1. Model A has a worse fit than any other model, with a single 𝜎^2^ value of 0.102. For context, this value means that the bounded Brownian motion process the data are fit to has a variance of 0.102 per million years (of log-transformed expression values). This is the average rate across all 14,556 genes and across all branches of the tree. In contrast to a single-rate model, both models B and C are significantly better fits to the data. Contrary to some hypotheses, both models find that SI lineages (𝜎^2^) have a lower rate of evolution than SC lineages (𝜎^2^; Table 1). There is also a difference between the models, with model C (the one in which *S. pennellii* LA0716 shares a rate with SI species) fitting significantly better. To further examine the evolution of *S. pennellii* LA0716, model D fits a three-parameter model, with this lineage assigned its own rate of evolution. This model is a significantly better fit than model C (*P*<0.00001; χ^2^ test with 1 degree of freedom), and demonstrates that *S. pennellii* LA0716 has a rate of evolution (𝜎^2^ in Table 1) that is slightly *lower* than SI species. This highly similar rate to SI species implies that it has only recently transitioned to self-compatibility, which is consistent with previous inferences about the timing of transition to SC in this particular accession (e.g. Rick and Tanksley 1981).

CAGEE also allows users to infer the number and direction of changes in gene expression levels along each branch of the tree. Figure 3 reports the number of genes that had “credible” increases and decreases in expression level under model D. Credible changes require that the credible intervals around states at parent and daughter nodes do not overlap, in order to account for uncertainty in our inferences. However, because of this, fewer credible changes will be inferred deeper in the tree, where credible intervals get wider. Therefore, while inferences about the identity of the genes changing along each branch is greatly strengthened by using credible changes (these genes are noted in the raw output from CAGEE), the absolute numbers of credible changes cannot be compared across branches, except for sister branches of equal length. For completeness, we show the total numbers of increases and decreases of gene expression in Supplementary Figure 4; as expected, these total numbers are more uniformly distributed across older and younger branches.

We assessed whether the genes identified as having credible increases or decreases in expression specifically on any SC branch (solid red branches in Figure 3) were significantly enriched for any biological process or molecular function gene ontology (GO) categories compared to genes with credible changes on any SI branch (black branches in Figure 3). This comparison specifically assesses gene expression evolution associated with a transition to SC, over and above “background” rates of expression evolution across the rest of the clade. Although fold enrichment was modest 1.20-1.36X; Supplementary Table 2), there were 11 terms significantly enriched (FDR<0.05) specifically on SC branches; these terms primarily focused on regulation of transcription, metabolic processes, and biosynthesis (Supplementary Table 2). Among the genes in these over-represented categories, a large fraction are transcription factors associated with development (e.g. WRKY and MADS Box), hormonal responses (including ethylene- and auxin-responsive transcription factors), and regulation of cell cycle (e.g. cyclins), in addition to protein kinases (Supplementary Table 2). This enrichment is consistent with increased expression changes in genes involved in cell division, differentiation, and development, that could follow transitions to SC.

## Discussion

Here, we have developed a new software package that enables the estimation of rates of gene expression evolution across a tree, CAGEE. Gene expression levels are much like many other continuous traits, and multiple papers have introduced phylogenetic comparative methods for studying gene expression (Bedford and Hartl 2009; Rohlfs et al. 2014; Rohlfs and Nielsen 2015; Catalán et al. 2019; Chen et al. 2019). However, as far as we are aware none of these methods allows genes to share evolutionary parameters, which precludes the analysis of genome-wide trends, either along the branches of a tree or between tissues/samples/conditions. To overcome this limitation, CAGEE calculates the likelihood of the data using the pruning algorithm (Felsenstein 1973) to facilitate the sharing of evolutionary parameters along branches of the species tree, providing more statistical power to test evolutionary hypotheses.

Fortunately, we were able to take advantage of much of the codebase of our existing software, CAFE (Hahn et al. 2005; De Bie et al. 2006; Hahn et al. 2007; Han et al. 2013; Mendes et al. 2020), which implements the pruning algorithm for the analysis of gene family sizes across a tree. While gene expression levels and gene family sizes differ in the type of data they represent (continuous vs. discrete) and their underlying evolutionary models (bounded Brownian motion vs. birth-death), many of the required likelihood calculations and software components are the same.

An important thing to consider for the input to CAGEE is the normalization used to make gene expression levels comparable across species. The data from wild tomatoes used here was normalized using TPM (transcripts per million; Wagner et al. 2012); other published datasets also use this normalization (Berthelot et al. 2018; Chen et al. 2019; El Taher et al. 2021). However, multiple other normalizations have also been used in comparative analyses, including RPKM (Brawand et al. 2011), FPKM (Catalán et al. 2019), and both TMM and CPM (Blake et al. 2020). Each normalization approach has its advantages and disadvantages, and we cannot yet strongly recommend one specific approach as input to CAGEE. The normalization method used will likely depend on the conditions under which samples are collected: if all species can be raised simultaneously in a greenhouse, vivarium, or growth chamber, we expect many fewer batch effects than in samples collected from the field, which will therefore necessitate different normalizations. However, even animals raised in a common environment—but fed different diets—can show many differences in gene expression not due to heritable change (e.g. Somel et al. 2008). Conversely, many between-sample normalization approaches (e.g. TMM, trimmed mean of M values; Robinson and Oshlack 2010) make the assumption that differences in gene expression between samples are rare. While such normalization is sensible in the context of testing for differential expression between samples from the same species, for a set of species that have been evolving independently for millions of years this is likely not an appropriate assumption.

CAGEE currently has multiple limitations, both in the available models that can be applied and in the types of data that can be analyzed. As mentioned earlier, many researchers have modeled gene expression using an OU process (Bedford and Hartl 2009; Rohlfs et al. 2014; Chen et al. 2019; Yang et al. 2019). Although OU models may be artifactually preferred over unbounded Brownian motion models due to a number of non-biological factors (see discussion in “New Approaches” above), it would still be helpful to be able to compare such a model to the bounded Brownian motion model used here. However, fitting such a model to genome-wide data is non-trivial: each gene must have its own mean expression value (μ), but possibly shared constraint parameters (α) across genes. We have the goal of implementing such a model in the near future, as well as other models commonly used in comparative methods research (e.g. Landis and Schraiber 2017; Boucher et al. 2018).

Beyond the evolutionary model applied to any dataset, there are multiple additional sources of variation that could be modeled. For instance, we have previously accounted for measurement error in a likelihood framework, using an empirically parameterized error model (Han et al. 2013). We can imagine both applying a similar model here to RNA-seq data, as well as extending CAGEE to more error-prone data such as single-cell sequencing. Such an extension would treat the level of expression in each cell within a cell type as an error-prone draw from an underlying distribution; one would then be able to infer the rate of evolution within and across cell-types across multiple species. The biggest obstacle to this approach may be in identifying homologous cell types across species (e.g. Tarashansky et al. 2021). In addition, not all genes necessarily share the same average rate of evolution; gamma-distributed rate categories can be used to model this variation among genes (cf. Ames et al. 2012; Mendes et al. 2020). As shown above, not accounting for this rate variation leads to a slight underestimate of 𝜎^2^, but also obscures interesting patterns of evolution among genes. Finally, the gene tree discordance found in many phylogenomic datasets implies that complex traits (such as expression levels) will also be controlled by discordant gene trees (Hahn and Nakhleh 2016; Hibbins and Hahn 2021). This underlying discordance can cause evolutionary rates to be overestimated (Mendes et al. 2018), and should be taken into account when seeking accurate parameter estimates (see discussion of wild tomato data below). Our goal is to include methods for dealing with all these sources of variation in future versions of CAGEE.

In terms of the types of data that can be analyzed, at present CAGEE is limited to positive, continuously varying traits (i.e. the BBM model). However, we also envision different ways to represent and model gene expression data, including as a ratio (e.g. male/female expression). Such a ratio, after log2-transformation, would be most appropriately modeled by an unbounded Brownian motion model since both negative and positive values are possible. This and other data types will be supported in future releases. Moreover, CAGEE does not have to analyze whole-genome or even molecular data: it can be applied to any single trait for which the BBM model is appropriate, even morphological traits. One intriguing application of CAGEE could be to suites of morphological traits that are hypothesized to share a common evolutionary rate parameter. If, for instance, there is a shift in body plan along some lineages, then multiple traits may all increase or decrease their rate of evolution at once, and CAGEE can be used to estimate these shared parameters. Even in the context of single-trait analyses, the pruning algorithm has been hailed as a solution for large-scale comparative analyses (Freckleton 2012). Importantly, the number of branches in a rooted, bifurcating tree with *n* tips is 2*n*-2, so that the number of calculations scales linearly with the number of species. This makes the pruning algorithm ideal for comparative datasets with large numbers of taxa (e.g. Hahn et al. 2005; FitzJohn 2012; Hiscott et al. 2016; Caetano and Harmon 2018; Mitov et al. 2020).

The analysis of data from a clade of wild tomatoes revealed a possibly unexpected result: the rate of ovule gene expression evolution among self-compatible (SC) species is twice as high as the rate among self-incompatible (SI) species (Table 1). This finding is contrary to some prior expectations—informed by research focused on male-female interactions, especially between interacting proteins in the reproductive tract (e.g. Swanson and Vacquier 2002; Clark et al. 2006)— that suggest that lineages might experience slower evolution after transitioning to self-compatibility. However, it is possible that global gene expression levels do not evolve in the same sort of tit-for-tat manner as interacting protein sequences, such that increases/decreases in male-expressed genes are not matched by increases/decreases in interacting female-expressed genes (or vice versa). Alternatively, only a very small subset of genes may evolve in this manner. Indeed, even prior studies comparing protein evolution have failed to find clear evidence of slower global evolutionary rates in more inbreeding species (e.g. Wong 2011). One caveat to the observed rate differences in our data is that underlying gene tree discordance, whether due to incomplete lineage sorting or introgression, can lead to artifactually higher rate estimates (Mendes et al. 2018; Hibbins and Hahn 2021). However, there is in fact less discordance among the SC lineages sampled here (Pease et al. 2016), which is the reverse of the pattern that would be required to explain our results.

If not due to underlying bias in our estimates, these findings still raise the question: why is ovule gene expression evolving more rapidly in SC than SI species? One possibility is that this increased rate is due to a relaxation of selection in SC species, possibly because genes involved in male-female interactions are no longer needed. If this were the case, we might expect to see a general decrease in expression levels in SC species; however, there appears to be no consistent directionality to the changes along SC branches (Figure 3, Supplementary Figure 4). Instead, an alternative hypothesis is that transitions to SC involve adaptation to new optima of ovule gene expression, compared to SI species that tend to maintain ancestral optima. For example, transitions to SC might favor greater investment in fewer ovules, because self-compatibility decreases the probability that each ovule within a flower will go unfertilized—an otherwise wasted investment under conditions (like SI) where receiving sufficient compatible pollen to fertilize each ovule is less predictable (Burd et al. 2009). The nature of these new optima might be even more complex, as traits like ovule size and number can vary with multiple reproductive and ecological conditions, and often trade-off with each other (Greenway and Harder 2007). Of the species examine here, for example, two SC lineages (*S. pimpinellifolium*, and *S. lycopersicon*—domesticated tomato) have significantly larger seeds than most of the SI lineages and SC *S. pennellii* (unpubl. data). Indeed, individual genes identified in our GO analysis are known to directly influence ovule and/or seed size in *Solanum* (e.g. *NOR-like1* [SOLYC07G063420.3.1; Han et al, 2014], *GRAS2* [SOLYC07G063940.2.1; Li et al. 2018], and *CRY2* [SOLYC09G090100.3.1; Fantini et al. 2019]). Some of our hypotheses could be evaluated with matching gene expression data from other (non-ovule) reproductive tissues. Analyses including pollen in the same SI and SC lineages, and/or data addressing alternative constraints and conditions shaping ovule evolution including ovule size and number (e.g. Mione and Anderson 1992), would be useful in teasing apart these hypotheses.

## Material and Methods

### Bounded Brownian motion model of expression evolution

The probability density of expression, 𝑝(𝑥, 𝑡), at time 𝑡 for evolutionary trajectories following a Brownian motion process starting at value 𝑥_𝑡0_ at time 𝑡_*_ is governed by the diffusion equation

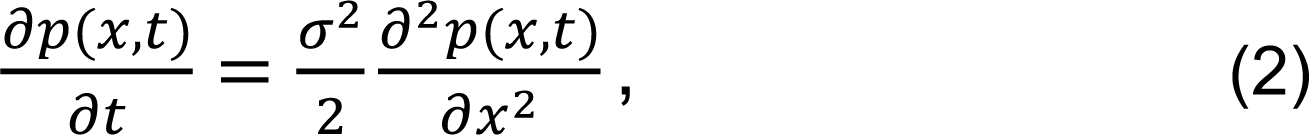

with initial condition 𝑝(𝑥, 𝑡_*_) = 𝛿(𝑥 − 𝑥_𝑡0_) where 𝛿 is the Dirac delta function. The reflective boundary condition at 𝑥 = 𝑥_m$n_ implies that the probability fluxes into and out of the boundary are balanced, imposing the boundary condition

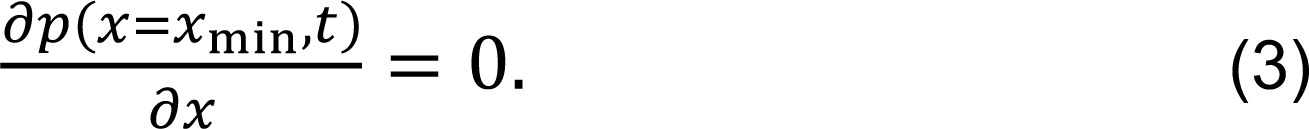

Note that 𝑝(𝑥, 𝑡) is identical to the transition density 𝑝(𝑥_𝑡_|𝑥_𝑡0_).

Without the reflecting boundary, 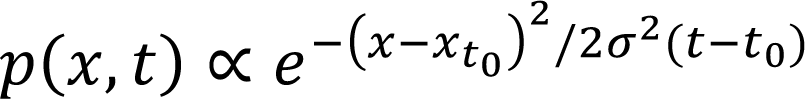 is a normal distribution with variance 𝜎^2^(𝑡 − 𝑡_*_). The variance therefore scales linearly with elapsed time, 𝑡 − 𝑡_*_. With the reflecting boundary, 𝑝(𝑥, 𝑡) is the sum of this spreading normal and its mirror image centered at 2𝑥_m$n_ − 𝑥_𝑡0_ . The analytical solution to this bounded process is helpful for understanding the behavior of 𝑝(𝑥, 𝑡), but is not used in CAGEE. In anticipation of implementing additional (and possibly more complicated) processes into CAGEE, we instead solve Eq. (2) numerically using the approach described in Boucher and Démery (2016). Briefly, the continuous diffusion equation is converted into a matrix equation by discretizing expression values into 𝑁 equal bins of width 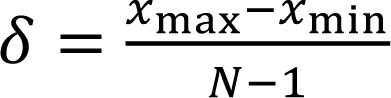. Following Boucher and Démery (2016), we have used a default *N*=200, but this number can be set by the user (see Results). This approach gives

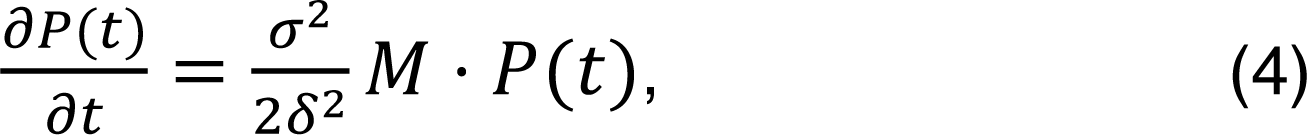

where 𝑃(𝑡) is the vector obtained by discretizing 𝑝(𝑥, 𝑡) and 𝑥_m78_ is the largest expression value accounted for. The matrix 𝑀 is tridiagonal with −2 on the diagonal except at the first and last diagonal entries which are −1. The sub- and supra-diagonal entries are 1. This equation has the matrix exponential solution

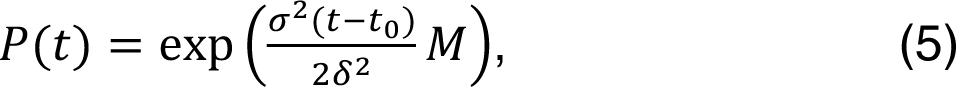

which is evaluated by diagonalizing 𝑀.

### Implementation of CAGEE

CAGEE is written in C++ and is compatible with the C++11 standard. A comprehensive manual and extensive unit tests facilitate further code development and maintenance. CAGEE is organized into modular components. A *clade* class, with references to a parent clade and any number of descendant clades, represents a tree structure, and a *gene_transcript* class represents the expression levels observed in the various species. These two classes comprise the fundamental data structures upon which CAGEE performs its analysis (Supplementary Figure 5).

Calculations are carried out by additional classes. The *optimizer* class has the responsibility of determining the 𝜎^2^ value with the highest likelihood, by comparing the likelihood of candidate values and searching the likelihood surface using the Nelder-Mead optimization algorithm. The work of computing the likelihood of a given 𝜎^2^ value is performed by a subclass of the *model* class, which for now is limited to a single *Base* model (allowing for further development in the future). After appropriate estimated values are found, the *transcript_reconstructor* class builds a possible set of transcript values for the entire tree (Supplementary Figure 5).

Performing the likelihood calculations requires extensive matrix operations; it is recommended (though not required) that these be passed off to a specialized library such as Intel’s MKL or Nvidia’s CUBLAS. If no external library is available, CAGEE will carry out these calculations (slowly) by itself. Creating the diffusion matrix (𝑀) requires calculation of eigenvalues and eigenvectors, and is computationally expensive. This work is performed by the Eigen linear algebra library (https://eigen.tuxfamily.org); various internal data structures also take advantage of Eigen classes. To enable faster searching, the matrix for an ancestral state vector of length 200 (the default in CAGEE) has been pre-computed and is included with CAGEE. Users who wish to use vectors of different lengths can specify this as an option.

Unit-testing is performed using the Doctest testing framework (https://github.com/doctest/doctest). At the time of writing more than 200 unit tests had been created, comprising more than 1200 individual assertions. For complex logging and debugging cases, CAGEE uses the EasyLogging framework (https://github.com/amrayn/easyloggingpp). C++ development is always made easier by using the Boost C++ libraries (https://www.boost.org/), so we include them as well in CAGEE.

### RNA-seq data from wild tomatoes

We briefly describe here the data collected from seven accessions of wild tomatoes (*S. lycopersicum* LA3475, *S. chmielewskii* LA1316, *S. pimpinellifolium* LA1589, *S. habrochaites* LA1777, *S. chilense* LA4117A, *S. pennellii* LA3778, and *S. pennellii* LA0716; all accession ID numbers from tgrc.ucdavis.edu). Further details are given in Moyle et al. (2021). Ovule RNA-seq was performed on between one to four (usually three) biological replicates (individual plants) from each accession. Plants were germinated from seed, and cultivated until flowering. For each replicate individual, ovules were dissected from mature, unpollinated flowers, flash frozen, and maintained at -80C until extraction. For each individual, all ovule collections were pooled into a single sample prior to library construction and sequencing on an Illumina HiSeq 2000.

Reads were mapped against the tomato reference genome (ITAG 2.4) and the number of reads mapped onto genic regions were estimated with featureCounts (Liao et al., 2014). We normalized the read counts from each library by calculating TPM (transcripts per million; Wagner et al. 2012) and then calculated the mean normalized read counts across all samples (individuals) within each accession. These means per accession were used as input to CAGEE.

To construct a species tree for use with CAGEE, we started with the topology given in Pease et al. (2016). Specifically, we used the tree found in the supplementary file Pease_etal_TomatoPhylo_RAxMLConcatTree_no1360_Fig2A.nwk, and pruned it to include only the accessions in our study using the software ETE (Huerta-Cepas et al. 2016). Using the “extend” method found in ETE, we converted this tree to ultrametric (same root-to-tip distance for all taxa). Setting the root age to 2.48 million years ago (following Pease et al. 2016), we were able to express all branches in millions of years. Analyses of GO enrichment were carried out using ShinyGO (Ge et al. 2020) with a false discovery rate of 0.05.

## Supplementary Material

Supplementary data are available at .

## Acknowledgements

We thank Mark Hibbins for assistance with the tomato phylogeny, Matthew Gibson for putting together the tomato gene expression data, and especially Dan Vanderpool for invaluable help in the initial development of CAGEE. Two reviewers provided helpful comments, and Scott Edwards pointed out relevant work that we had previously missed. This work was supported by National Science Foundation grants DEB-1856469 to L.C.M. and DBI-2146866 to M.W.H.

## Data Availability

Raw reads for each sample library are available at NCBI BioProject PRJNA714065. The CAGEE software is available at https://github.com/hahnlab/CAGEE.

## Figures and Tables

## Supplementary Figures

**Supplementary Figure 1.**
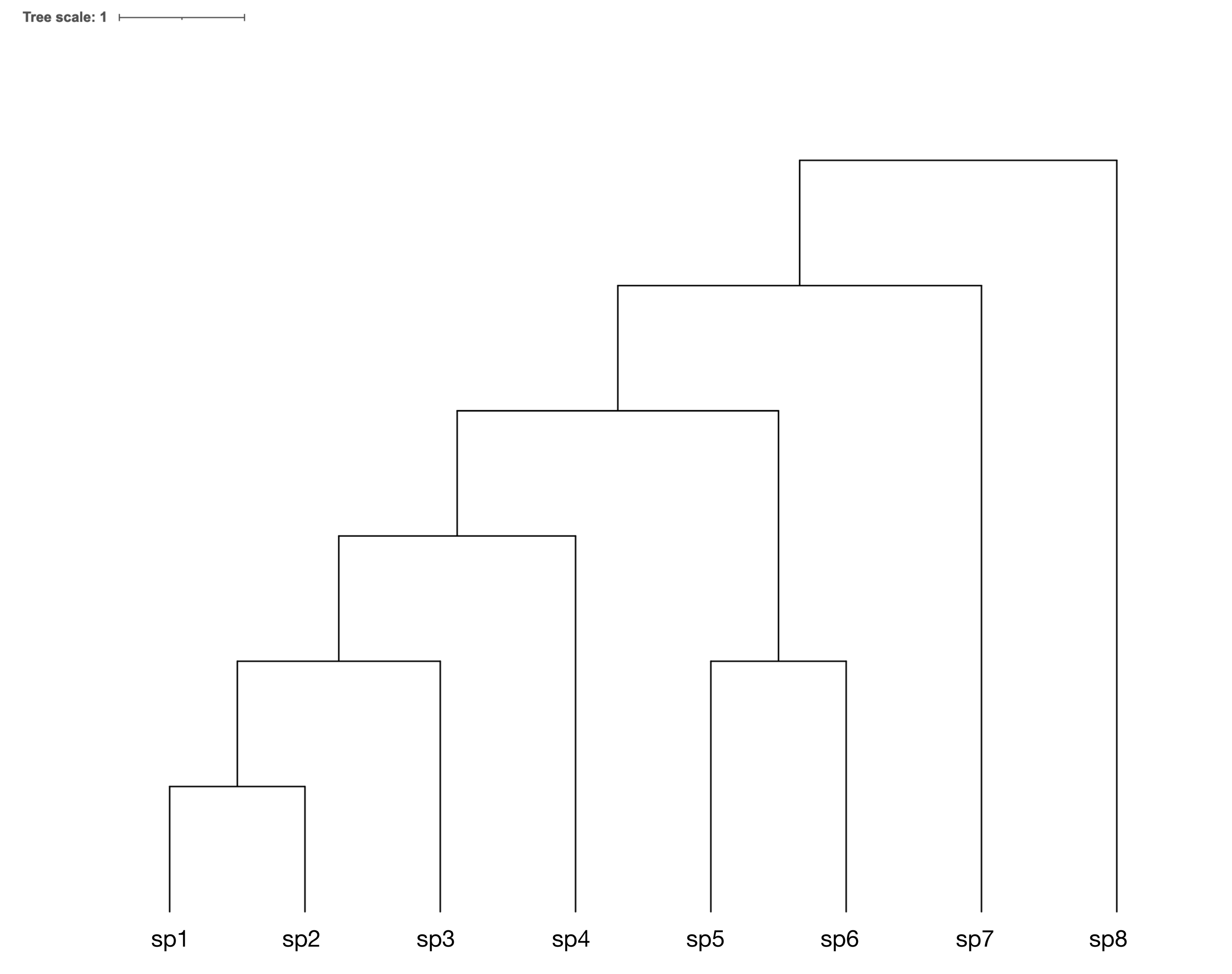
The tree used for simulations. The Newick-formatted tree string with branch lengths is: ((((sp1:1,sp2:1):1,sp3:2):1,sp4:3):1,((sp5:2,sp6:2):2)):1,sp7:5):1,sp8:6)

**Supplementary Figure 2.**
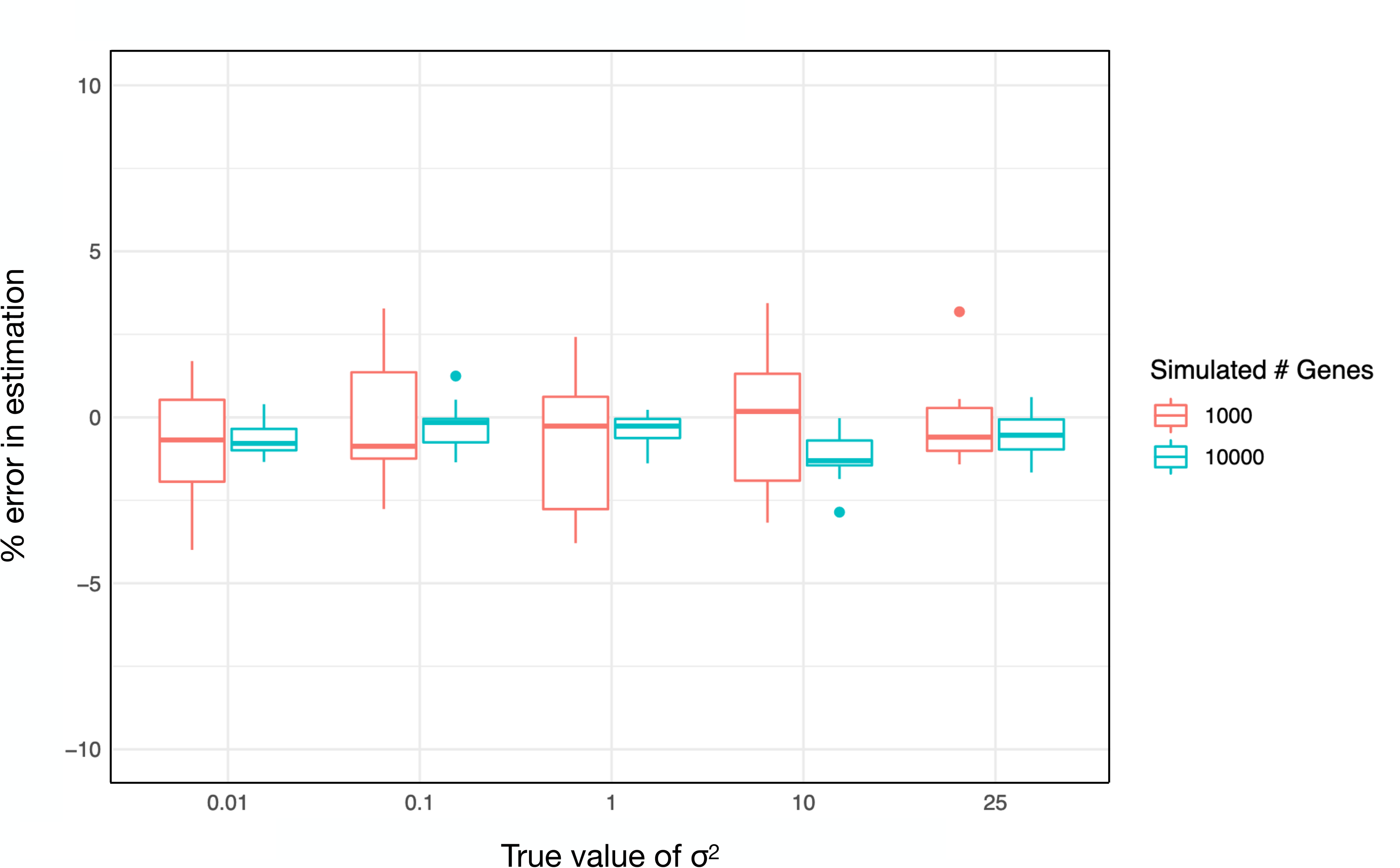
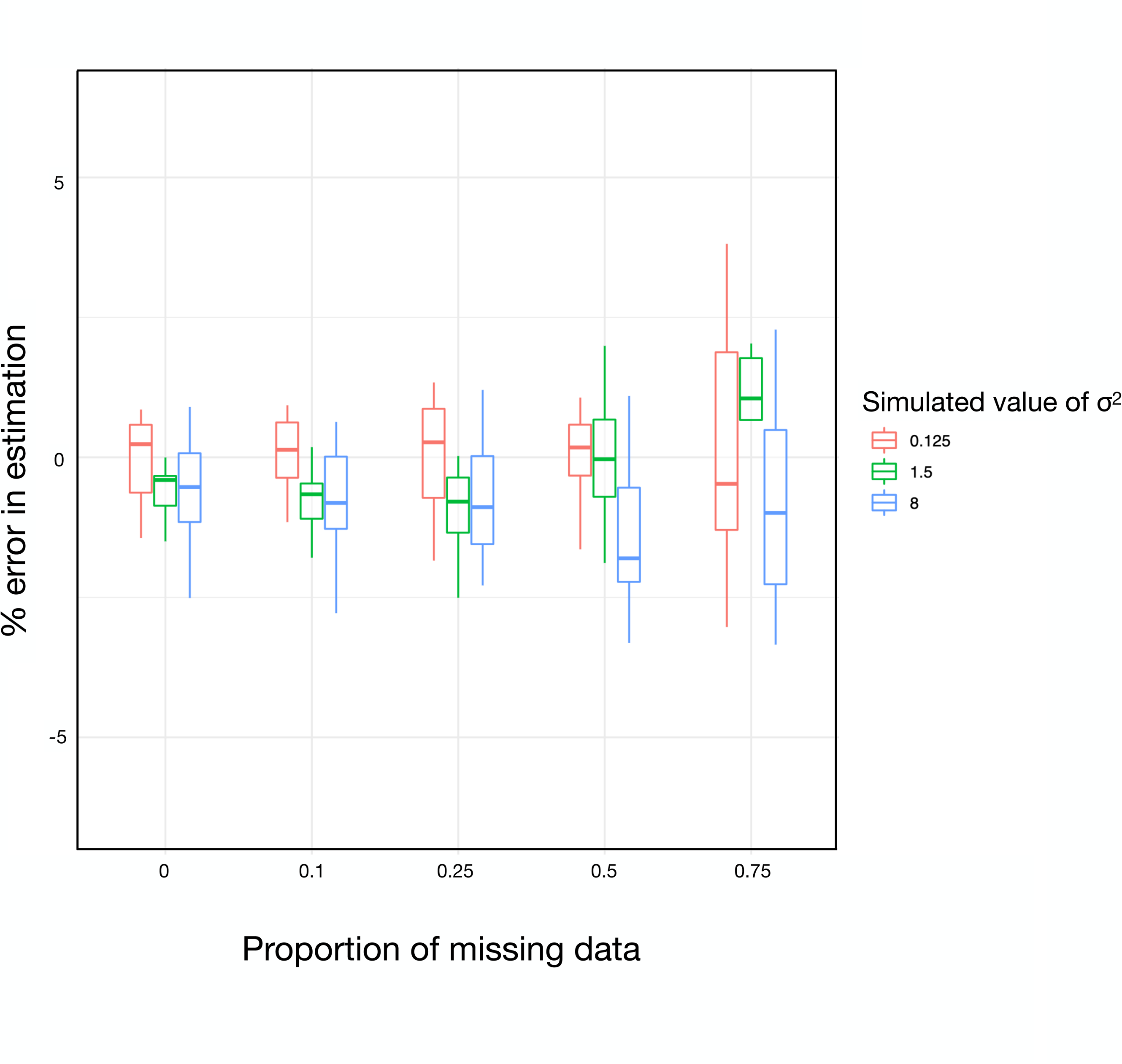
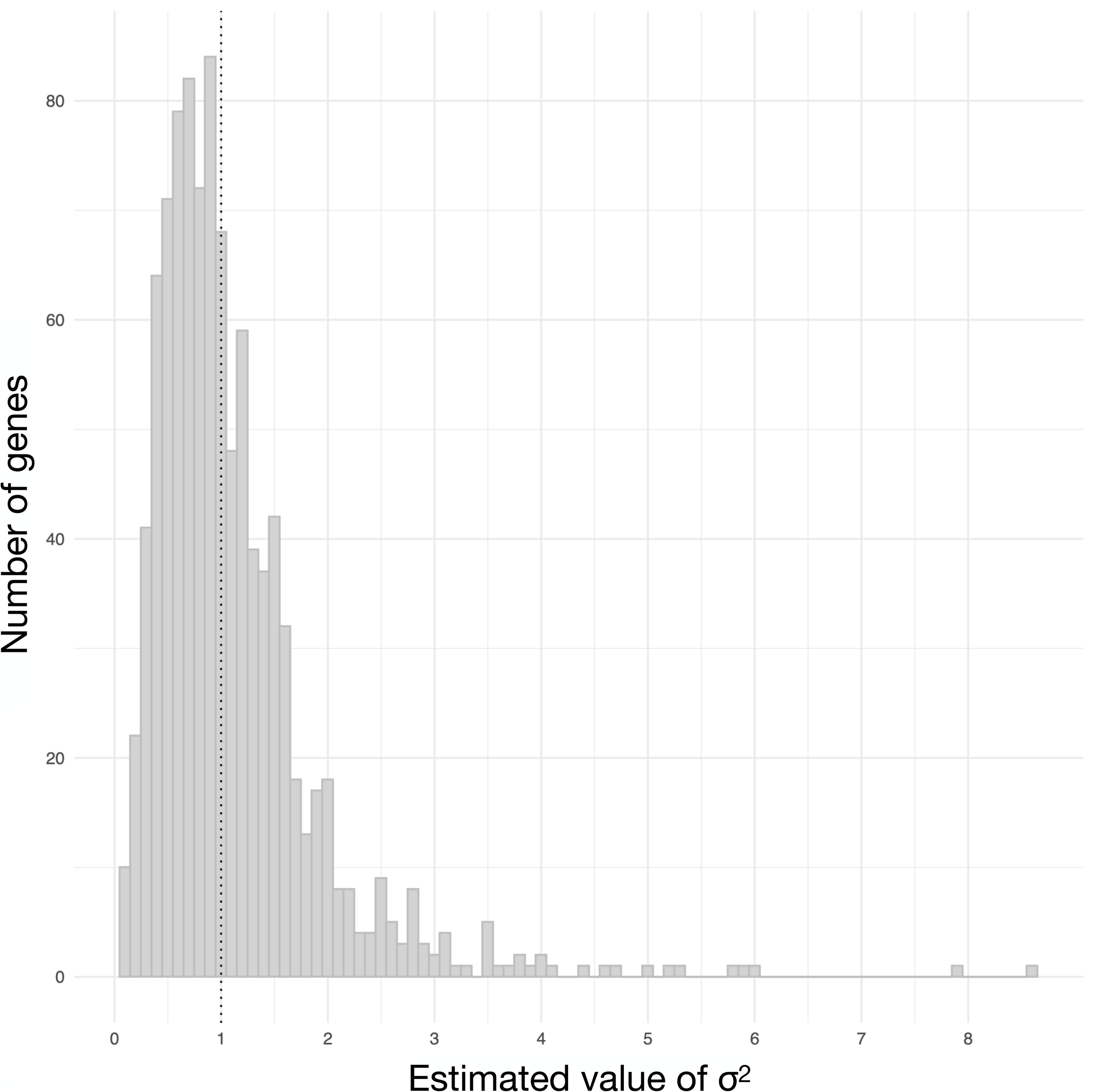
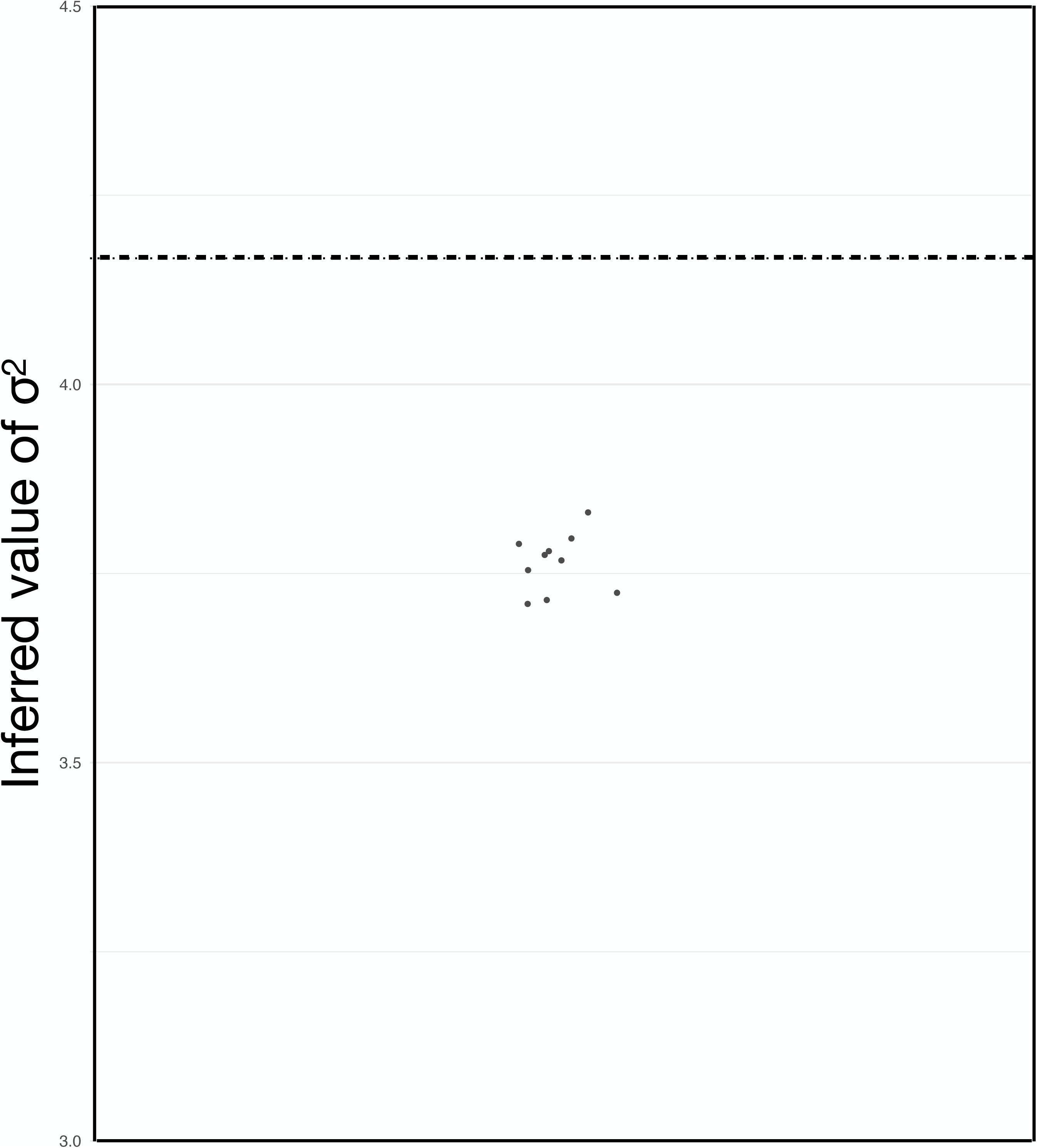
Accuracy of CAGEE. A) This figure is the same as Figure 2 in the main text, but the ancestral state vector has length *N*=500 (Figure 2 uses *N*=200). B) For each of three different simulated values of 𝜎^2^, we randomly removed different amounts of data from an input dataset with 1,000 genes (the tree is the same as in all other simulations). C) For 1,000 genes simulated with 𝜎^2^=1 (dashed vertical line), we ran CAGEE independently on each one to estimate 𝜎^2^. D) We combined three datasets of 1,000 genes each simulated with three different values of 𝜎^2^ (we repeated these simulations 10 times). The 10 estimates of 𝜎^2^ on the combined datasets were slightly downwardly biased compared to the expected value (dashed horizontal line). Each dot represents each of the 10 estimates, with jitter added for clarity,

**Supplementary Figure 3.**
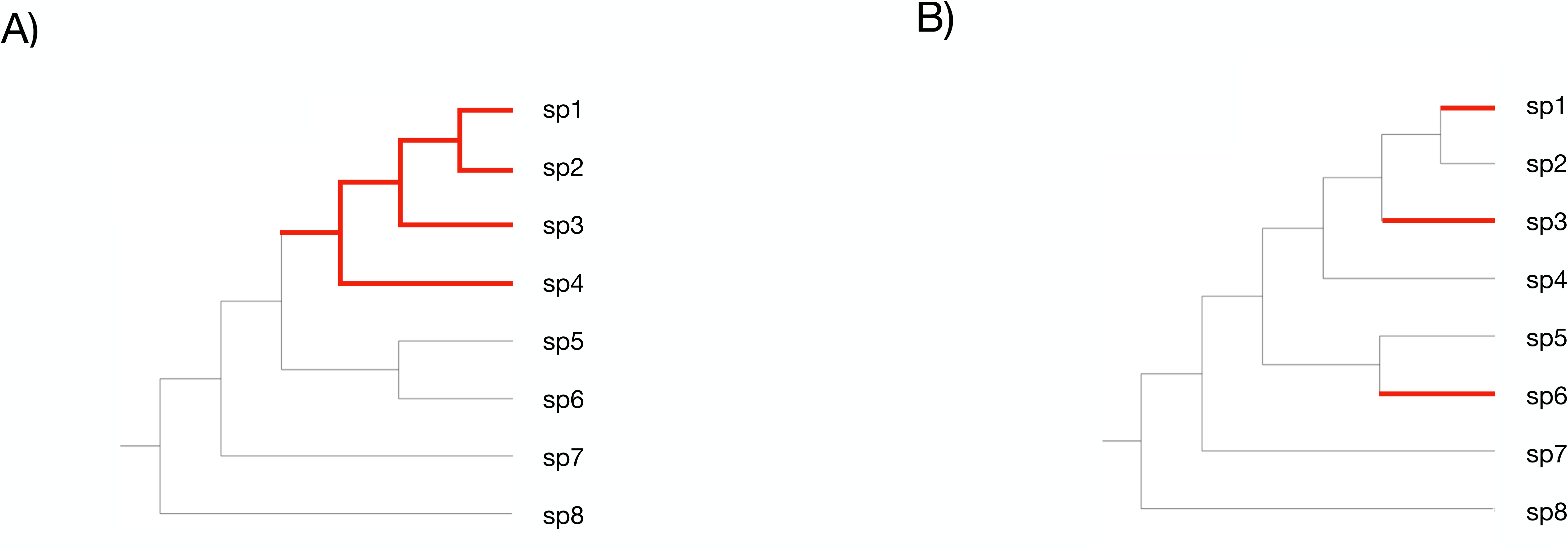
Trees used for simulations with lineage-specific values of 𝜎^2^. All black branches share a rate parameter (𝜎^2^_1_), and all red branches share a rate parameter (𝜎^2^_2_). This “sigma_tree” is specified in CAGEE with the Newick string: ((((sp1:2,sp2:2):2,sp3:2):2,sp4:2):2,((sp5:1,sp6:1):1)):1,sp7:1):1,sp8:1) A) All black branches share a rate parameter (𝜎^2^_1_), and all red branches share a rate parameter (𝜎^2^_2_). This “sigma_tree” is specified in CAGEE with the Newick string: ((((sp1:2,sp2:1):1,sp3:2):1,sp4:1):1,((sp5:1,sp6:2):1)):1,sp7:1):1,sp8:1)

**Supplementary Figure 4.**
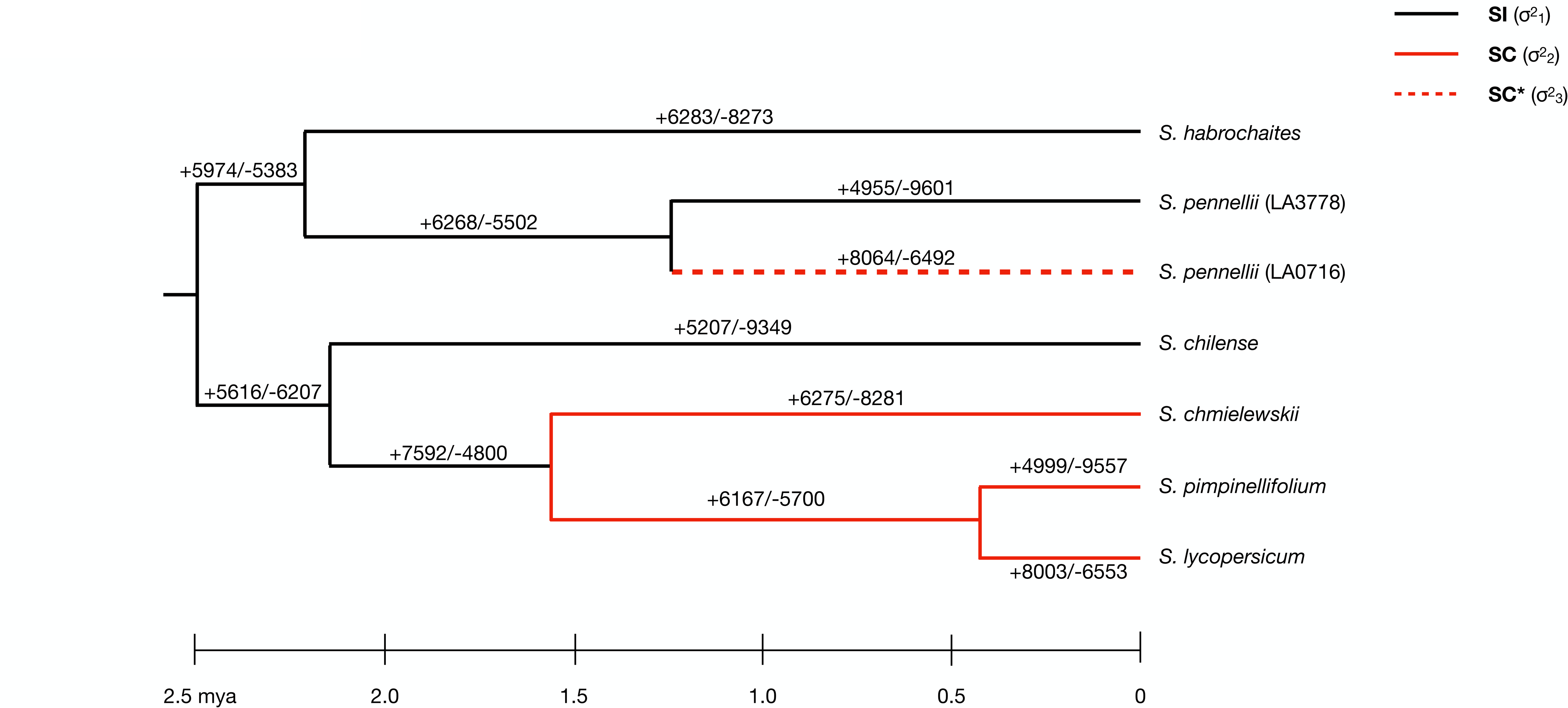
Changes in gene expression along the tomato phylogeny. This figure is the same as Figure 3 in the main text, but all increases and decreases are reported, regardless of whether they are “credible”.

**Supplementary Figure 5.**
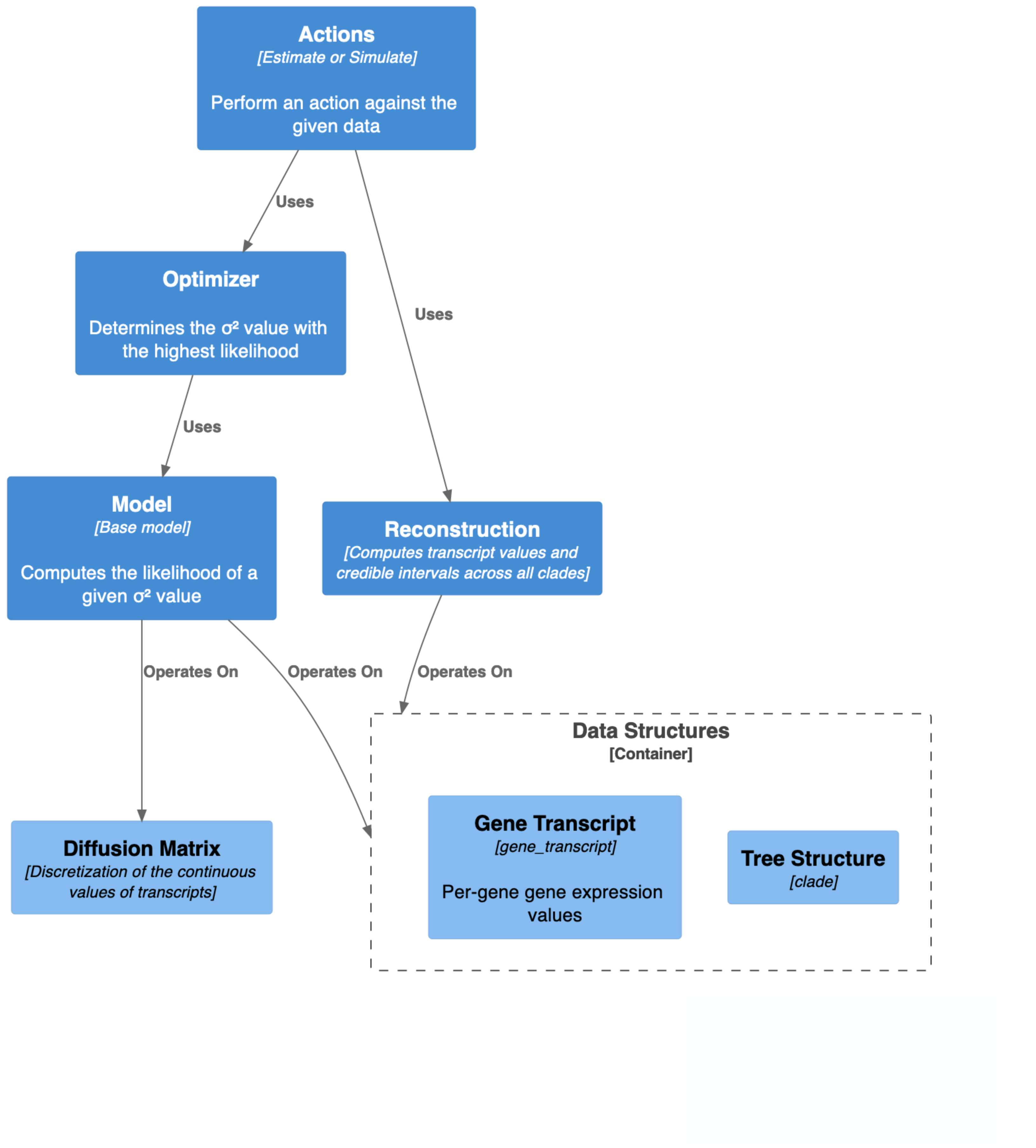
Component diagram for the CAGEE software.

**Supplementary Table 1.**
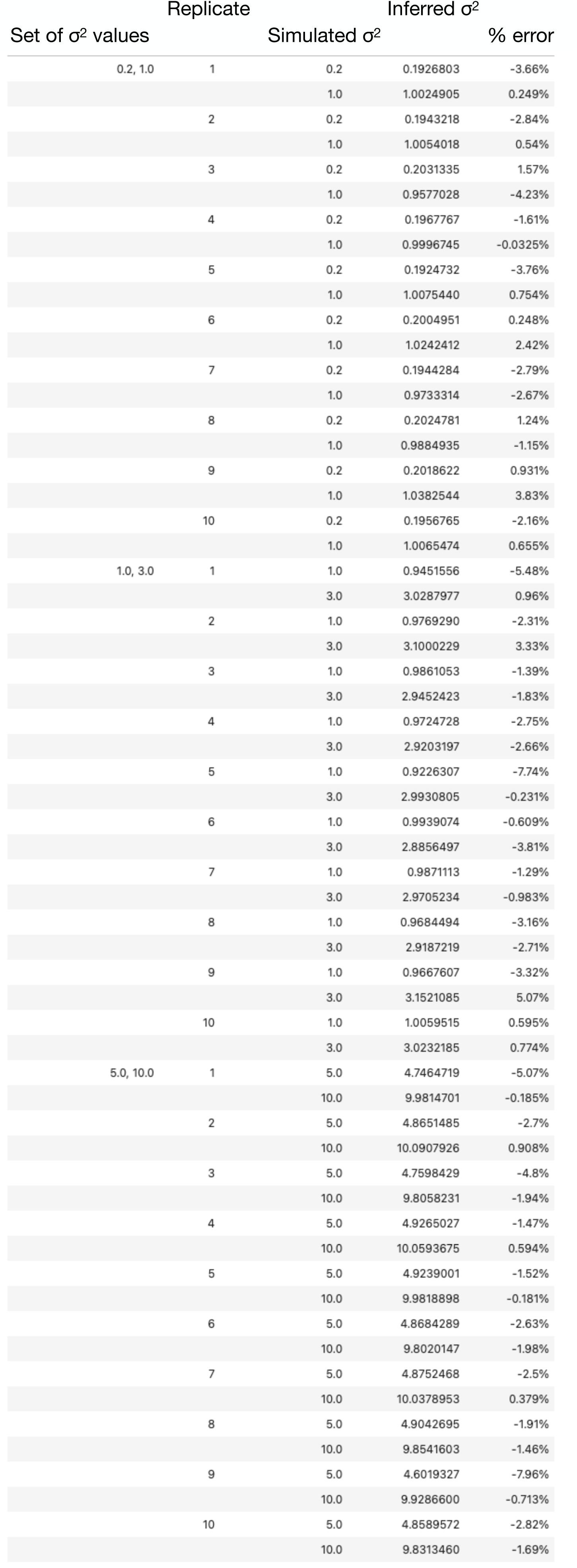

